# Induction of long-term hyperexcitability by memory-related cAMP signaling in isolated nociceptor cell bodies

**DOI:** 10.1101/2024.07.13.603393

**Authors:** Alexis Bavencoffe, Michael Y. Zhu, Sanjay V. Neerukonda, Kayla N. Johnson, Carmen W. Dessauer, Edgar T. Walters

**Author notes:** Corresponding author: Dr. Edgar T. Walters, Department of Integrative Biology and Pharmacology McGovern Medical School at UTHealth, Houston, Texas 77030. These authors contributed equally to this work (Alexis Bavencoffe, Michael Y. Zhu). email addresses: Alexis Bavencoffe, Michael Zhu, Sanjay V. Neerukonda, Kayla N. Johnson, Carmen W. Dessauer, Edgar T. Walters.

## Abstract

Persistent hyperactivity of nociceptors is known to contribute significantly to long-lasting sensitization and ongoing pain in many clinical conditions. It is often assumed that nociceptor hyperactivity is mainly driven by continuing stimulation from inflammatory mediators. We have tested an additional possibility: that persistent increases in excitability promoting hyperactivity can be induced by a prototypical cellular signaling pathway long known to induce late-phase long-term potentiation (LTP) of synapses in brain regions involved in memory formation. This cAMP-PKA-CREB-gene transcription-protein synthesis pathway was tested using whole-cell current clamp methods on small dissociated sensory neurons (primarily nociceptors) from dorsal root ganglia (DRGs) excised from previously uninjured (“naïve”) rats. Six-hour treatment with the specific Gαs-coupled 5-HT4 receptor agonist, prucalopride, or with the adenylyl cyclase activator, forskolin, induced long-term hyperexcitability (LTH) in DRG neurons that manifested 12-24 hours later as action potential (AP) discharge (ongoing activity, OA) during artificial depolarization to -45 mV, a membrane potential that is normally subthreshold for AP generation. Prucalopride treatment also induced significant long-lasting depolarization of resting membrane potential (from -69 to -66 mV), enhanced depolarizing spontaneous fluctuations (DSFs) of membrane potential, and indications of reduced AP threshold and rheobase. LTH was prevented by co-treatment of prucalopride with inhibitors of PKA, CREB, gene transcription, and protein synthesis. As in the induction of synaptic memory, many other cellular signals are likely to be involved. However, the discovery that this prototypical memory induction pathway can induce nociceptor LTH, along with reports that cAMP signaling and CREB activity in DRGs can induce hyperalgesic priming, suggest that early, temporary, cAMP-induced transcriptional and translational mechanisms can induce nociceptor LTH that might last for long periods. An interesting possibility is that these mechanisms can also be reactivated by re-exposure to inflammatory mediators such as serotonin during subsequent challenges to bodily integrity, “reconsolidating” the cellular memory and thereby extending the duration of persistent nociceptor hyperexcitability.

**Highlights:**

- Nociceptor long-term hyperexcitability (LTH) can be induced by a 5-HT4R agonist.
- 5-HT4R-induced LTH manifests as ongoing activity during modest depolarization.
- Enhanced ongoing activity is associated with long-term potentiation of DSFs.
- Induction of LTH depends upon PKA, CREB, transcription, and protein synthesis.
- Nociceptor LTH may be triggered by conserved memory-related plasticity mechanisms.

## 1. Introduction

Much is known about mechanisms that trigger acute pain or maintain chronic states of painful hypersensitivity (Basbaum et al., 2009; Finnerup et al., 2021), far less about how temporary noxious or inflammatory events can induce persistent painful states (Price and Ray, 2019). Although many mechanisms, including non-neuronal mechanisms, are likely to be involved (e.g., Li et al., 2019), a plausible hypothesis is that highly conserved mechanisms of memory induction found in invertebrate nervous systems (Kandel, 2001; Davis, 2023) and the mammalian brain (Nguyen and Woo, 2003) are important for inducing persistent amplification of neural function in pain pathways. This general hypothesis (Walters, 1991; Woolf and Walters, 1991; Ji et al., 2003) was strengthened by the discovery that long-term synaptic potentiation (LTP) induced by activation of N-methyl-D-aspartate (NMDA)-sensitive glutamate receptors is prominent not only in mammalian and invertebrate synapses involved in memory (Huang and Stevens, 1998) but also in synapses within multiple components of pain pathways (Li et al., 2019), including primary nociceptor synapses (Sandkühler, 2007). At many of these synapses, the induction of persistent potentiation involves an early increase in cAMP synthesis (Nguyen and Woo, 2003). Furthermore, studies in several species revealed a prototypical (but simplified) intracellular signaling pathway for inducing synaptic memory in which cyclic AMP (cAMP) activates protein kinase A (PKA), which activates the transcription factor cAMP response element binding protein (CREB), which induces gene transcription that ultimately enhances the synthesis of plasticity-related proteins (Silva et al., 1998; Kandel, 2012).

cAMP signaling can also contribute to the induction of late-phase LTP at C-fiber synapses (Liu and Zhou, 2015), to induction of long-term hyperexcitability (LTH) in rat cortical neurons (Cudmore and Turrigiano, 2004) and *Aplysia* nociceptors (Scholz and Byrne, 1988), and to induction of persistently enhanced responsiveness of rodent nociceptors to inflammatory mediators (hyperalgesic priming) (Ferrari et al., 2015). These observations, and the importance of persistent hyperexcitability in nociceptors for driving persistent pain states (Walters et al., 2023), led us to test the hypothesis that this prototypical plasticity induction pathway can induce LTH in rat nociceptors that substantially outlasts the inducing stimulus. To limit the numerous potential factors that may contribute to LTH induction, we have adopted a simple in vitro approach (Fig. 1A), similar to that originally used to define this induction pathway at synapses in dissociated *Aplysia* neurons (Kandel, 2001). Like those studies, we use an agonist of Gαs-coupled serotonin (5-HT) receptors as the inducing agent. To avoid complicating effects from other 5-HT receptors, rather than 5-HT we use the specific Gαs-coupled 5- HT4 receptor agonist, prucalopride, which we demonstrated previously to stimulate acute PKA- dependent hyperexcitability in dissociated nonpeptidergic nociceptors from rats (Lopez et al., 2021).

**Figure 1.**
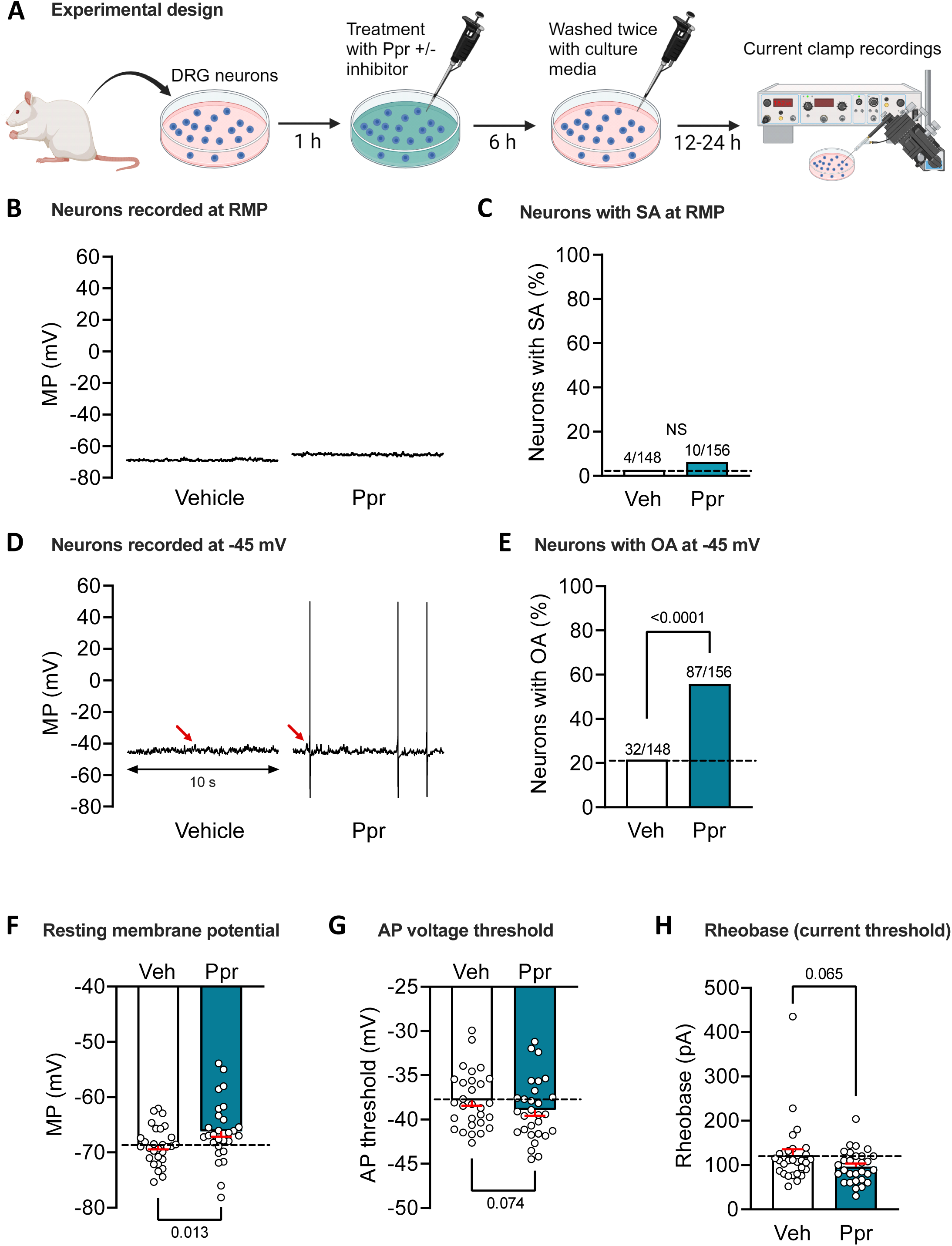
5-HT4 receptor agonist prucalopride (Ppr) induces long-term hyperexcitability (LTH) in small (≤ 30 µm soma diameter) sensory neurons. ***A***, Experimental design (created with BioRender.com). ***B-E***, Long-term effects on excitability of treatment with Ppr (1 µM) versus vehicle (0.01 % DMSO). Representative 10-s recordings at RMP ***B***), or held under current clamp at -45 mV (***D***). Corresponding proportions of neurons exhibiting any AP discharge at RMP (SA), ***C***) or at -45 mV (OA), ***E***). Above each bar is the number of neurons exhibiting discharge over the total number sampled. Comparisons using Fisher’s exact test with *p* values reported above each bar. Red arrows indicate the largest subthreshold depolarizing spontaneous fluctuation (DSFs) in each trace. ***F-H***, Impact of Ppr 6-hour treatment on other measures of excitability, RMP (***F***), AP voltage threshold (***G***), and rheobase (***H***). Datasets are graphed as mean ± SEM. Each circle represents the mean value for that condition per day of recording when both vehicle and prucalopride were tested. Comparisons used paired t-tests with *p* values reported in each panel. *p*<0.05 was considered significant. AP, action potential; DRG, dorsal root ganglion; MP, membrane potential; N.S., non-significant; OA, ongoing activity; RMP, resting membrane potential; SA, spontaneous activity; Veh, vehicle.

Here we show that this prototypical synaptic memory induction pathway can also induce LTH in nociceptor somata. While many transcriptional, translational, and post-translational alterations of nociceptors have been linked to persistent pain (Reichling and Levine, 2009; Ferrari et al., 2015; Price and Ray, 2019; Gale et al., 2022; Ghosh and Pan, 2022), this is the first demonstration of an early induction pathway for nociceptor hyperexcitability that is both transcription- and translation-dependent.

## 2. Materials and Methods

### 2.1. Animals

All procedures were in accordance with the guidelines of the International Association for the Study of Pain and were approved by the McGovern Medical School Animal Care and Use Committees. Male Sprague-Dawley rats were used, 8-9 weeks old, 250-300g (Envigo, USA). Possible sex differences will be the subject of a separate study. Rats were allowed to acclimate to the vivarium for at least four days before beginning experiments.

### 2.2. Dissociation and culture of dorsal root ganglion (DRG) neurons

Euthanasia was performed with an intraperitoneal injection of pentobarbital/phenytoin solution (Euthasol, 0.9 ml, Virbac AH, Inc.) followed by transcardial perfusion of ice-cold phosphate buffered saline (PBS, Sigma-Aldrich). DRGs were exposed from the dorsal side of the spinal column, excised, and transferred to high-glucose DMEM culture medium (Sigma-Aldrich) containing trypsin TRL (0.3 mg/ml, Worthington Biochemical Corporation) and collagenase D (1.4 mg/ml, Roche Life Science). After 40 min of enzymatic digestion at 34°C under constant shaking, DRGs were washed with DMEM by two successive centrifugations and triturated with two fire-polished glass Pasteur pipettes of decreasing diameters. The resulting mixed culture was then plated on 8-mm glass coverslips (Warner Instruments) coated with poly-L-ornithine 0.01 % (Sigma-Aldrich) in DMEM without serum or growth factors, and incubated overnight at 37°C, 5% CO_2_ and 95% humidity.

### 2.3. Whole-cell electrophysiology

All patch-clamp recordings were performed 18-30 h after dissociation (12-24 h after experimental treatment) at room temperature (∼ 21°C) with a Multiclamp 700B amplifier (Molecular Devices). DRG mixed cultures were observed at 20x magnification on IX-71 (Olympus, Japan) inverted microscope and recorded in extracellular recording solution (ECS) containing (in mM): 140 NaCl, 3 KCl, 1.8 CaCl_2_, 2 MgCl_2_, 10 HEPES, and 10 glucose, adjusted to pH 7.4 with NaOH and 320 mOsM with sucrose. Borosilicate glass (Sutter Instrument) patch pipettes were obtained using a horizontal P-97 puller (Sutter Instrument) and fire-polished with a MF-830 microforge (Narishige) to a final pipette resistance of 3-8 MΩ when filled with an intracellular solution containing (in mM): 134 KCl, 1.6 MgCl_2_, 13.2 NaCl, 3 EGTA, 9 HEPES, 4 Mg-ATP, and 0.3 Na-GTP, adjusted to pH 7.2 with KOH and 300 mOsM with sucrose. Whole cell recordings were conducted on sensory neuron somata having diameters ≤ 30 µm. After securing a tight seal (>3 GΩ), the plasma membrane was ruptured to reach whole-cell configuration under voltage clamp at -60 mV. Recordings were acquired with Clampex v10.7 (Molecular Devices). The liquid junction potential was calculated to be ∼4.3 mV and not corrected, meaning the actual potentials may have been ∼4.3 mV more negative than indicated in the measurements presented herein.

### 2.4. Excitability tests

Resting membrane potential (RMP) was recorded for at least 60 s after the membrane potential stabilized, along with any spontaneous activity (SA). Neurons with RMP less negative than - 40 mV under our recording conditions were excluded. Membrane potential was then held at -60 mV manually (± 2 mV) under current clamp for action potential (AP) voltage threshold and rheobase measurements, as well as classification of nonaccommodating (NA) and rapidly accommodating (RA) DRG neurons (Odem et al., 2018). This was done with an incrementing series of 2-s depolarizing current injections (5-20 pA steps), stopping at 2x rheobase. Neurons exhibiting ≥ 2 action potentials (APs) during any step were classified as NA, while neurons showing a single AP at the beginning of the step were classified as RA. AP threshold was defined conservatively as the most depolarized subthreshold membrane potential during the 2-s steps (up to 1 step above the step where discharge occurred). Membrane potential was then held at -45 ± 2 mV under current clamp for 30 s. The occurrence of any APs during this moderately depolarized period was defined as ongoing activity (OA) (Odem et al., 2018).

### 2.5. Quantification of depolarizing spontaneous fluctuations (DSFs)

Measurement of DSFs from whole-cell current clamp recordings utilized the Frequency- Independent Biological Signal Identification (FIBSI) program. FIBSI was written using the Anaconda (v2019.7.0.0, Anaconda, Inc) distribution of Python (v3.5.2) with dependencies on the NumPy and matplotlib.pyplot libraries [CASSIDY 2020]. The updated version of FIBSI used in this study is an automated program that incorporates the algorithm used to analyze DSFs in our prior publications (Odem et al., 2018; Berkey et al., 2020; Garza Carbajal et al., 2020; Laumet et al., 2020; Bavencoffe et al., 2022; North et al., 2022; Bavencoffe et al., 2024; Tian et al., 2024). DSFs were detected in 30-s recordings sampled at 10 kHz with Multiclamp 700B (Molecular Devices) and filtered with a 10 kHz Bessel filter, using a Ramer-Douglas-Peucker algorithm. Estimation of the resting membrane potential (RMP) at each point uses a 1-s sliding median function. For each point, the program then reports the amplitudes and durations of DSFs and hyperpolarizing spontaneous fluctuations (HSFs). Cutoff values for DSFs and HSFs were set at minimum amplitudes and durations of 1.5 mV and 5 ms, respectively. Amplitudes for the suprathreshold DSFs eliciting APs were estimated conservatively as the difference of the most depolarized potential reached by the largest subthreshold DSF within the recording from the sliding median at that point. This value was substituted for each AP (Odem et al., 2018; Bavencoffe et al., 2022). A minimal interval of 200 ms between any two APs was required for the second AP to be substituted by the maximal suprathreshold DSF amplitude in the recording period. If the interval between two APs was less than 200 msec, then a substitution was performed only if the peak of a separate DSF or HSF was detected between the APs (Odem et al., 2018). The original FIBSI source code and detailed tutorial are available for free use, modification, and distribution on a Github (GitHub, Inc.) repository titled “FIBSI Project” by user “rmcassidy” (https://github.com/rmcassidy/FIBSI_program).

### 2.6. Drugs

All pharmacological agents used in this study were purchased from Sigma-Aldrich and reconstituted in DMSO. Adenylyl cyclase activator forskolin, 5-HT4 receptor agonist prucalopride, 5- HT4 receptor antagonist GR113808, and PKA inhibitor H-89 stock solutions were prepared at 10 mM, protein synthesis inhibitor cycloheximide at 20 mM, CREB inhibitor 666-15 at 5 mM, and transcription inhibitor actinomycin D at 1 mg/ml. LTH was induced by 6 h incubation with prucalopride (Ppr, 1 µM) or forskolin (Fsk, 1 µM) in DMEM culture medium, followed by two washes with DMEM.

### 2.7. General experimental design and statistical analysis

To address our hypothesis that a cAMP-PKA-CREB-transcription-translation pathway is sufficient to induce nociceptor LTH, our general design was to treat mixed dissociated lumbar and thoracic DRG cultures for 6 h with prucalopride +/- inhibitors predicted to at least partially block the hypothesized pathway, testing for effects on somal excitability 12-24 h later with electrophysiological test protocols under whole-cell current clamp (Fig. 1A). Contributions of individual components of the tested signaling pathway were assessed by incubating neurons with prucalopride plus a pharmacological inhibitor of the selected component. In each inhibitor study, we took advantage of our ability to run several experimental groups with multiple neurons sampled from a single rat (i.e., an individual experiment). This allowed us to decrease variability across experiments by excluding experiments in which LTH criteria were not met for acceptable positive and negative controls. In each experiment on neurons from a single rat that tested the effects of an inhibitor, 4 groups were included, each with 3-8 neurons: negative control (vehicle), positive control (prucalopride, Ppr), inhibitor alone, and Ppr + inhibitor groups. To answer the question of whether an inhibitor prevented the induction of LTH by Ppr (entirely or partially) we compared the proportion of sampled neurons with OA at -45 mV in the Ppr group (positive control) to the proportion in the Ppr + inhibitor group. The criterion for inclusion of an experiment in the study was conservative evidence (based on our previous studies) for LTH induced in the positive controls (defined as OA in ≥ 40% of neurons in the Ppr group) and lack of LTH induced in the negative controls (defined as OA in ≤ 40% of neurons in the vehicle group). If either criterion was not met, the experiment was excluded and no data from that rat was used in the inhibitor study. The inhibitor-alone group was used to see if any inhibitors had persistent effects on excitability independent of Ppr. All reported data are presented as medians, means ± SEM, or proportion of neurons expressed as percentage of neurons sampled. Normal distribution of populations was assessed with the Shapiro–Wilk test. For normally distributed datasets, comparisons were done by one-way ANOVA or Brown-Forsythe and Welch ANOVA (when standard deviations were unequal) followed by Dunnett’s multiple comparisons tests using Prism version 8.3 (GraphPad Software). All other datasets were analyzed with the Kruskal–Wallis test followed by Dunn’s multiple comparison tests. Comparisons between two datasets were made with paired or unpaired t-tests for normally distributed data, and Mann-Whitney U tests for other data. Comparisons of proportions were made using Fisher’s exact test with Bonferroni corrections for multiple comparisons. Detailed statistical results are reported in the figures and their legends. Statistical significance was considered as p < 0.05, except after Bonferroni corrections for multiple comparisons of proportions, as stated in the legends.

## 3. Results

### 3.1. Exposure to 5-HT4 receptor agonist prucalopride for several hours induces long-term hyperexcitability (LTH)

Our use of the term LTH is based on studies in *Aplysia* sensory neurons (Walters and Ambron, 1995; Liao et al., 1999; Weragoda et al., 2004; Weragoda and Walters, 2007; Mihail et al., 2019) that defined LTH as hyperexcitability persisting for approximately 1 d or more after an induction event, which was based on similar terminology from in vitro studies of long-term facilitation of *Aplysia* sensory synapses (Kandel, 2001) and late, protein-synthesis dependent LTP of hippocampal synapses (Abraham et al., 1991; Abbas et al., 2015). Here we asked whether nociceptor LTH lasting 12-24 h can be induced in vitro by treatment of dissociated small DRG neurons (diameter ≤ 30 µm, primarily TRPV1-expressing nociceptors) (Bedi et al., 2010; Wu et al., 2013; Odem et al., 2018) for several hours with a 5-HT receptor agonist. We used prucalopride (Ppr, 1 µM), a specific agonist of Gαs-coupled 5-HT4 receptors, which we found previously to induce acute hyperexcitability in dissociated nociceptors (Lopez et al., 2021). Limited preliminary studies (not shown) revealed no obvious LTH after treating cultures with Ppr for 2 or 4 h, so we used 6-h treatments in all experiments (Fig. 1A). Compared to treatment with vehicle (0.01% DMSO in DMEM), treatment with Ppr for 6 h beginning 1 h after plating induced LTH manifested as repetitive discharge (ongoing activity, OA) of APs in putative nociceptors 12-24 h after washout. While no significant increase was found in neurons exhibiting OA at RMP (spontaneous activity, SA) (Fig. 1B, C), the proportion of neurons with OA when the membrane potential was artificially depolarized for 30 s to -45 mV was significantly elevated by Ppr treatment (Fig. 1D, E). This OA at -45 mV was accompanied by significant depolarization of RMP (Fig. 1F), and trends for hyperpolarization of AP voltage threshold (Fig. 1G), and reduction of AP current threshold (rheobase) (Fig. 1H) compared with the corresponding effects of vehicle treatment. These results show that temporary exposure of nociceptor somata to a 5-HT agonist known to stimulate Gαs-coupled 5-HT4 receptors and to activate protein kinase A (PKA) in dissociated nociceptors (Lopez et al., 2021) induces LTH lasting at least 12-24 h after the stimulation period.

### 3.2. LTH represents a persisting aftereffect of cAMP signaling

Induction of nociceptor LTH by 6-h treatment with Ppr indicated that activation of adenylyl cyclase (AC) and consequent cAMP signaling is sufficient to induce hyperexcitability persisting for about 1 day after treatment. To further investigate whether the persisting hyperexcitability involves a memory-like aftereffect of cAMP signaling, we asked whether it a) requires persistent activation of the Gαs-coupled 5-HT4 receptors and b) can also be induced by 6-h treatment with a direct activator of AC. Ppr is a potent stimulator of 5-HT4 receptor-induced PKA activation in dissociated nociceptors (K_i_ ∼5 nM) (Lopez et al., 2021), so we asked whether continuing exposure to extremely low concentrations of Ppr that might persist overnight (long after the washes in the DMEM solution and after replacement of the DMEM) could continue to stimulate the nociceptors. This unlikely possibility was refuted by our finding that when the same 6-h Ppr treatment was followed immediately by overnight incubation with a potent, specific 5-HT4 receptor antagonist, GR113808 (1 µM), significantly enhanced incidence of OA at -45 mV still occurred, which was not significantly different from the enhanced OA incidence induced by Ppr in the absence of a 5-HT4 receptor antagonist following the Ppr treatment (Fig. 2A, B). This is consistent with Ppr activating AC to induce a persisting, memory-like effect. We did not observe any significant effects of treatment with Ppr followed by GR113808 or with GR113808 alone on other measures of excitability (RMP, AP voltage threshold or rheobase) compared with vehicle treatment (0.01% DMSO), while incubation with Ppr alone produced significant hyperpolarization of AP voltage threshold (Table 1).

**Figure 2.**
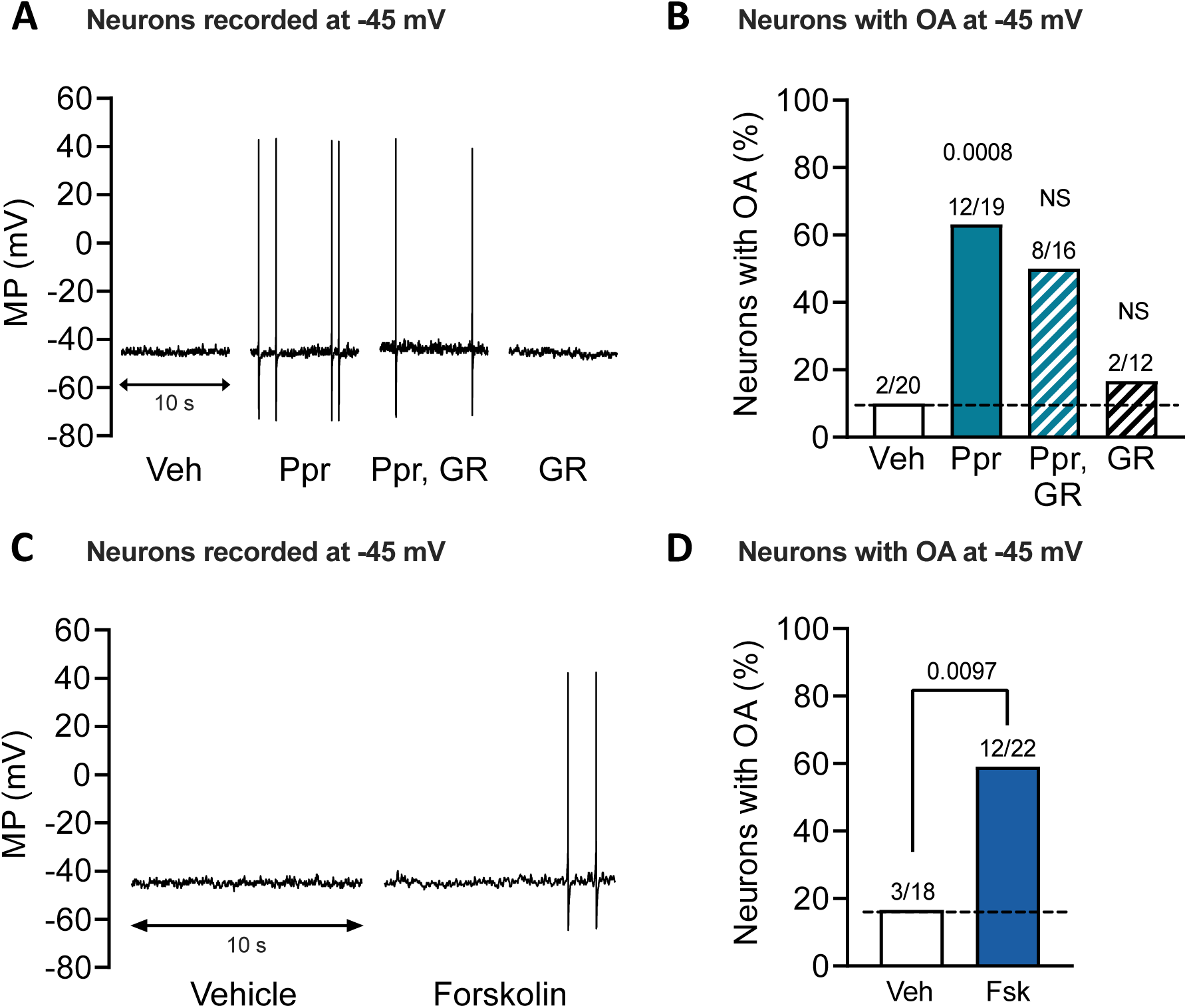
LTH is not attributable to continuing stimulation by Ppr and can be induced by direct stimulation of adenylyl cyclase. ***A***, 10-s representative traces of neurons recorded when clamped at - 45 mV after prior 6-h treatment with vehicle (“Veh”, 0.01% DMSO), Ppr (1 µM) alone, Ppr followed by overnight treatment with 5-HT4 receptor blocker GR113808 (1 µM), or GR113808 alone (“GR”). ***B***, proportions of neurons with OA at -45 mV in each experimental condition. ***C***, 10-s representative traces of neurons recorded at -45 mV after prior 6-h treatment with vehicle (0.01% DMSO) or adenylyl cyclase activator forskolin (1 µM). D, proportions of neurons with OA at -45 mV. For ***B*** and ***D***, the number of neurons exhibiting OA over the total number sampled is reported above each bar. Comparisons performed against the vehicle condition with Fisher’s exact test. In ***B***, *p*<0.017 was considered significant after Bonferroni correction for 3 comparisons while in ***D***, *p*<0.05 was considered significant for a single comparison. Fsk, forskolin; MP, membrane potential; NS, non-significant; OA, ongoing activity; Ppr, prucalopride; Veh, vehicle.

**Table 1:** Effects of prucalopride and inhibitors of 5-HT4 receptor, PKA, CREB, gene transcription, and protein synthesis on electrophysiological measures of excitability in NA neurons isolated from naïve rats.

| Inhibitor tested (target) | Testing condition | RMP (mV)<sup>a</sup> | AP volt. thresh. (mV)<sup>a</sup> | Rheobase (pA)<sup>a</sup> | C<sub>m</sub> (pF)<sup>a</sup> | SA incidence (%)<sup>b</sup> |
| --- | --- | --- | --- | --- | --- | --- |
| GR113808 (5-HT<sub>4</sub>R) Delayed<sup>c</sup> | Vehicle | -66.8<br>(-80.3, -59) (20) | -35<br>(-46.7, -24.9) (20) | 120<br>(60, 240) (20) | 30.5<br>(18, 43) (20) | 0<br>(0/20) |
|  | Ppr | -66.4<br>(-76.2, -48.7) (19) | -39.2<br>(-50, -23.8) (17) * | 100<br>(60, 220) (17) | 33<br>(21, 40) (18) | 5.3<br>(1/19) |
|  | Ppr, GR | -63.9<br>(-76.7, -52.1) (16) | -36.7<br>(-48.4, -27.8) (16) | 100<br>(40, 220) (16) | 33.5<br>(23, 55) (16) | 0<br>(0/16) |
|  | GR | -61.3<br>(-80.1, -56.2) (12) | -37.3<br>(-47, -30.1) (12) | 80<br>(20, 200) (11) | 27<br>(14, 39) (12) | 8.3<br>(1/12) |
| H-89 (PKA) | Vehicle | -68.5<br>(-72.9, -58.3) (15) | -37.6<br>(-47.8, -24.9) (15) | 100<br>(60, 240) (15) | 30<br>(18, 46) (15) | 0<br>(0/15) |
|  | Ppr | -67.2<br>(-88.7, -51.7) (15) | -39.2<br>(-51.3, -23.8) (15) | 80<br>(0, 220) (15) | 33<br>(20, 40) (15) | 0<br>(0/15) |
|  | Ppr + H-89 | -63.7<br>(-73.6, -50) (16) | -37.9<br>(-46.3, -29.1) (16) | 100<br>(40, 180) (16) | 27.5<br>(19, 45) (16) | 0<br>(0/16) |
|  | H-89 | -63.2<br>(-73, -48.1) (14) | -34.7<br>(-47.8, -27.6) (14) | 90<br>(20, 240) (14) | 27<br>(15, 33) (14) | 7.1<br>(1/14) |
| 666-15 (CREB) | Vehicle | -71.5<br>(-80.6, -60.3) (27) | -38.9<br>(-54.4, -29) (27) | 120<br>(20, 600) (27) | 31<br>(20, 42) (27) | 7.4<br>(2/27) |
|  | Ppr | -69.5<br>(-81.1, -45) (26) | -40.6<br>(-47.8, -32.2) (26) | 90<br>(20, 280) (26) | 32.5<br>(18, 41) (26) | 11.5<br>(3/26) |
|  | Ppr + 666-15 | -67.2<br>(-75.7, -50.6) (33) ** | -38<br>(-47.3, -31.7) (31) | 90<br>(20, 340) (32) | 31<br>(15, 44) (33) | 0<br>(0/33) |
|  | 666-15 | -70.7<br>(-81.8, -53.1) (16) | -40.9<br>(-47.9, -29) (15) | 100<br>(20, 220) (16) | 36<br>(26, 43) (16) | 0<br>(0/16) |
| ActD | Vehicle | -71.2<br>(-79.3, -53.6) (20) | -40.3<br>(-44.5, -29.4) (20) | 90<br>(20, 220) (20) | 31.5<br>(15, 41) (20) | 0<br>(0/20) |
| (mRNA synthesis) | Ppr | -71.3<br>(-77.9, -53.7) (22) | -41.1<br>(-47.6, -34.3) (22) | 80<br>(20, 140) (22) | 29<br>(17, 39) (21) | 4.5<br>(1/22) |
| | Ppr + ActD | -75.3<br>(-83.9, -68.8) (13) * | -33<br>(-41.3, -28.1) (13) \$\$\$ | 180<br>(20, 340) (13) # | 29<br>(16, 39) (13) | 0<br>(0/13) |
| | ActD | -75.2<br>(-79.3, -58.2) (12) | -29.4<br>(-35.5, -22.9) (12) \$\$\$ | 240<br>(20, 360) (12) ### | 29.5<br>(18, 38) (12) | 8.3<br>(1/12) |
| Chx<br>(protein synthesis) | Vehicle | -71.9<br>(-77.7, -61.1) (15) | -38.4<br>(-43.8, -29.4) (15) | 100<br>(20, 240) (15) | 31<br>(21, 40) (15) | 0<br>(0/15) |
|  | Ppr | -71.4<br>(-81, -54.3) (17) | -41<br>(-48.8, -36.4) (17) | 60<br>(20, 180) (17) | 29<br>(14, 45) (17) | 11.8<br>(2/17) |
| | Ppr + Chx | -71.1<br>(-79.6, -63.7) (21) | -38<br>(-49.7, -29.8) (20) | 140<br>(40, 240) (20) | 25.5<br>(17, 37) (20) \$ | 0<br>(0/21) |
|  | Chx | -72.2<br>(-83.6, -59.9) (9) | -39.5<br>(-44.1, -30.9) (9) | 120<br>(40, 180) (9) | 32<br>(21, 34) (9) | 0<br>(0/9) |

<sup>a</sup> Values are reported as medians (minimum, maximum) (number of cells sampled). <sup>b</sup> Values are reported as incidences in percentages (number of cells showing spontaneous firing / number of cells sampled). <sup>c</sup> GR113808 application was delayed until after the washout of Ppr, and it remained overnight. All other inhibitors were applied concurrently with Ppr. Each experimental group was tested against vehicle. Statistical tests: Kruskal-Wallis followed by Dunn's multiple comparisons tests (\* p<0.05, \*\* p<0.01, \*\*\* p<0.0001); Brown-Forsythe and Welch ANOVA followed by Dunnett's T3 multiple comparisons tests (# p<0.05, ## p<0.01, ### p<0.001); ordinary one-way ANOVA followed by Dunnett's multiple comparisons tests (\$ p<0.05, \$\$ p<0.01, \$\$\$ p<0.001). ActD, actinomycin D; AP volt. thresh., action potential voltage threshold; Chx, cycloheximide; C<sub>m</sub>, membrane capacitance; Ppr, prucalopride; RMP, resting membrane potential; SA, spontaneous activity.

To confirm that AC activation is sufficient to induce LTH, we incubated the cultures for 6 h with forskolin (1 µM) the day before excitability testing. Forskolin-induced LTH was revealed by a significant increase in the proportion of neurons exhibiting OA when held at -45 mV (Fig. 2C, D) compared with vehicle controls (0.01% DMSO). Unlike Ppr-induced LTH, forskolin-induced LTH was not accompanied by significant alterations of RMP, AP threshold, DSF amplitudes, or other electrophysiological properties (Table 2), although a trend for more frequent large (>5 mV) DSFs was found in neurons recorded at -45 mV the day after forskolin treatment (*p* = 0.064, vehicle versus forskolin, Mann–Whitney U test). The difference between Ppr treatment and forskolin treatment might have been because the sample sizes for forskolin were substantially smaller than for Ppr, and possibly because ACs other than those coupled to 5-HT4 receptors may produce additional effects that complicate the electrophysiological aftereffects induced by 5-HT4-coupled ACs.

**Table 2:**
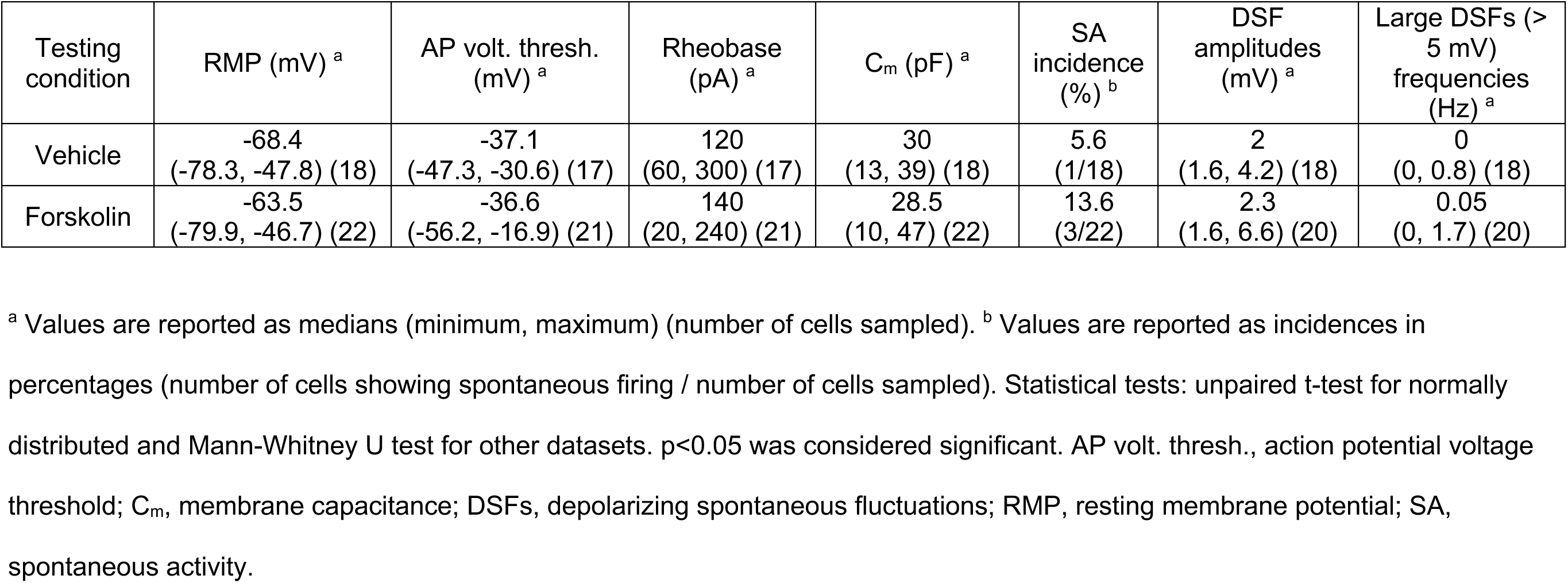
Effects of adenylyl cyclase activator forskolin on electrophysiological measures of excitability in NA neurons isolated from naïve rats.

| Testing condition | RMP (mV) <sup>a</sup> | AP volt. thresh. (mV) <sup>a</sup> | Rheobase (pA) <sup>a</sup> | C<sub>m</sub> (pF) <sup>a</sup> | SA incidence (%) <sup>b</sup> | DSF amplitudes (mV) <sup>a</sup> | Large DSFs (> 5 mV) frequencies (Hz) <sup>a</sup> |
| --- | --- | --- | --- | --- | --- | --- | --- |
| Vehicle | -68.4<br>(-78.3, -47.8) (18) | -37.1<br>(-47.3, -30.6) (17) | 120<br>(60, 300) (17) | 30<br>(13, 39) (18) | 5.6<br>(1/18) | 2<br>(1.6, 4.2) (18) | 0<br>(0, 0.8) (18) |
| Forskolin | -63.5<br>(-79.9, -46.7) (22) | -36.6<br>(-56.2, -16.9) (21) | 140<br>(20, 240) (21) | 28.5<br>(10, 47) (22) | 13.6<br>(3/22) | 2.3<br>(1.6, 6.6) (20) | 0.05<br>(0, 1.7) (20) |

<sup>a</sup> Values are reported as medians (minimum, maximum) (number of cells sampled). <sup>b</sup> Values are reported as incidences in percentages (number of cells showing spontaneous firing / number of cells sampled). Statistical tests: unpaired t-test for normally distributed and Mann-Whitney U test for other datasets. $p < 0.05$ was considered significant. AP volt. thresh., action potential voltage threshold; C<sub>m</sub>, membrane capacitance; DSFs, depolarizing spontaneous fluctuations; RMP, resting membrane potential; SA, spontaneous activity.

### 3.3. Prucalopride-induced LTH is prevented by inhibitors of PKA, CREB, gene transcription, and protein synthesis

Our core hypothesis about a fundamental cAMP-activated memory induction pathway being sufficient to induce nociceptor LTH was tested with the powerful pharmacological strategy described in section 2.7. This leveraged our ability to test 4 groups with multiple DRG neurons in each group in individual rats (each rat representing a single experiment in each inhibitor study). By requiring that each accepted experiment have at least 3 neurons in each of 4 groups (negative control [vehicle, DMSO 0.02 or 0.11 %, depending upon the drug used in the same experiment]), positive control [prucalopride, Ppr], inhibitor alone, and Ppr + inhibitor), we decreased variability within each inhibitor study by excluding experiments in which minimal LTH criteria were not met for acceptable positive and negative controls (see section 2.7). The procedures were identical to those presented in Figure 1 except that 2 additional groups of 3-8 neurons were added in each experiment: 1 group of neurons was treated for 6 h with one of the following inhibitors H-89 (10 µM) for PKA, 666-15 (0.5 µM) for CREB, actinomycin D (ActD, 1 µg/ml) for gene transcription, and cycloheximide (Chx 20 µM) for protein synthesis; another group in each experiment was treated with Ppr combined with one of the inhibitors just listed (Fig. 3A). Each inhibitor study included at least 3 experiments with 4 groups per experiment and 9-26 neurons sampled per group. Co-application of Ppr with the listed inhibitor of PKA (Fig. 3B), CREB (Fig. 3C), gene transcription (Fig. 3D), or protein synthesis (Fig. 3E) during the induction period significantly decreased the proportion of neurons exhibiting OA the next day in neurons artificially depolarized to -45 mV. No significant effects were found on the incidence of OA at -45 mV when each inhibitor was applied alone. Significant depression of other manifestations of hyperexcitability by the inhibitors (Table 1) included depolarization of RMP when combining Ppr with either 666-15 or actinomycin D, a reduction in AP voltage threshold and rheobase with actinomycin D alone or when combined with Ppr, and a reduction in the membrane capacitances of the sampled neurons when exposed to Ppr supplemented with cycloheximide when compared to vehicle (Table 1). We did not observe significant effects of Ppr when compared with vehicle on other measures of excitability (Table 1). This probably reflects the reduced statistical power when samples sizes are reduced by inclusion of neurons solely from individual experiments in which all 4 conditions were tested (vehicle, Ppr, Ppr + inhibitor, inhibitor). In sum, the consistent effects of the inhibitors on the induction of persistent OA confirms our hypothesis that LTH induced in small sensory neurons by Ppr exposure depends upon a PKA-CREB-gene transcription-protein synthesis signaling pathway.

**Figure 3.**
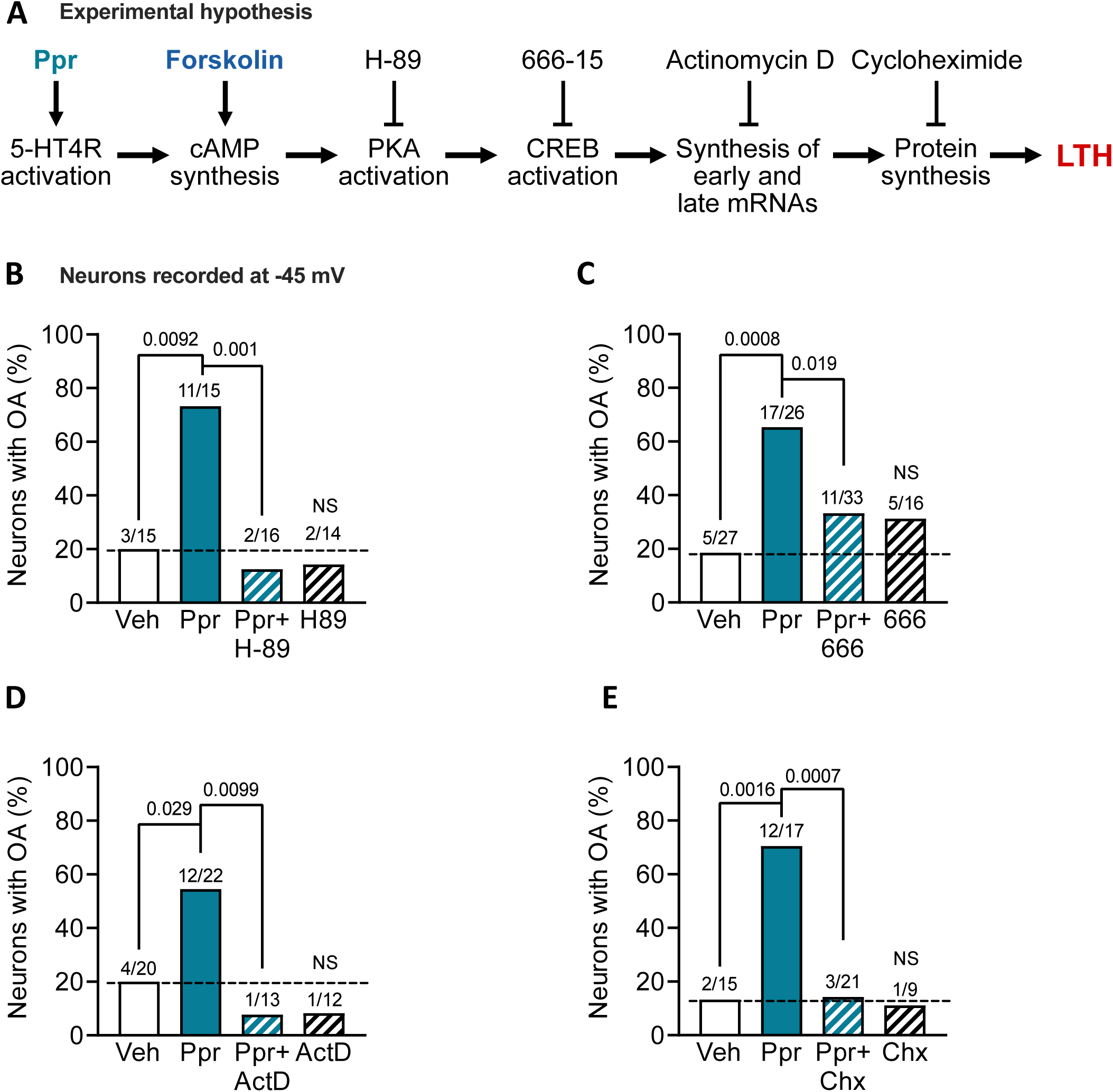
Induction of LTH depends upon PKA, CREB, gene transcription, and protein synthesis. ***A***, diagram of the experimental hypothesis with the inhibitors and activators employed for testing. ***B-E***, proportions of neurons with OA at -45 mV in vehicle (0.02 or 0.11 % DMSO, depending upon the inhibitor used), the day after 6-h treatment with Ppr (1 µM) alone or in combination (“Ppr + X”) with an inhibitor of PKA (H-89, 10 µM, ***B***), CREB (666-15, 0.5 µM, ***C***), gene transcription (actinomycin D, ActD, 1 µg/ml, ***D***) or protein synthesis (cycloheximide, Chx, 20 µM, ***E***), and each inhibitor alone. The number of neurons exhibiting OA over the total number sampled is reported above each bar. Comparisons with Fisher’s exact tests followed by Bonferroni correction for multiple comparisons. For comparisons of OA proportions between Ppr against the vehicle or Ppr + inhibitor, the significance level was * p< 0.025, ** p<0.005, and *** *p*<0.0005 for two comparisons. For vehicle against inhibitor alone, *p*< 0.05 was considered significant. LTH, long-term hyperexcitability; NS, non-significant; OA, ongoing activity.

### 3.4. Prucalopride-induced LTH is associated with potentiation of depolarizing spontaneous fluctuations of membrane potential (DSFs)

In principle, OA in dissociated nociceptors held at -45 mV can only be generated by a reduction in AP voltage threshold and/or an increase in the frequency of large DSFs that bridge the gap between the holding potential and AP threshold (Odem et al., 2018; Bavencoffe et al., 2024). LTH after Ppr treatment was associated with a modest decrease in AP threshold (Fig.1E), which can account for at least part of the increased OA. Automated analysis of DSFs at the -45 mV holding potential in each neuron in the inhibitor studies shown in Figure 3 revealed that Ppr treatment also induced long-term potentiation of DSFs (see Fig 4A for an enlargement of the Ppr recording in Fig 1D to highlight the DSFs). This potentiation was expressed as two significant effects: larger DSF amplitudes (Fig. 4B), and higher frequency in each neuron of larger DSFs (≥ 5 mV) that would be likely to approach AP threshold (Fig. 4C). Both potentiation effects were prevented by co-incubation of Ppr with actinomycin D to inhibit gene expression (Fig. 4B, C). Inhibition of PKA with H89 and inhibition of protein synthesis with cycloheximide in Ppr-treated neurons significantly reduced the frequency of larger (≥ 5 mV) DSFs (Fig. 4C) while a trend toward a reduction in DSF amplitude was observed in neurons treated with Ppr co-incubated with cycloheximide (Fig. 4B). Inhibition of CREB with 666-15 had no significant effect on any of the DSF measures (Fig. 4B-C). When given alone for 6 h, the inhibitors had no statistically significant long-term effects on the DSF measures (Fig. 4B-C), except for actinomycin D, which significantly reduced DSF amplitudes below the level of the DMSO controls. These results indicate that potentiated DSFs, in addition to lowered AP voltage thresholds, drive the OA at -45 mV during LTH induced by Ppr treatment, and they suggest that long-term potentiation of the DSFs involves induction mechanisms that depend upon PKA activity, gene transcription, and protein synthesis.

**Figure 4.**
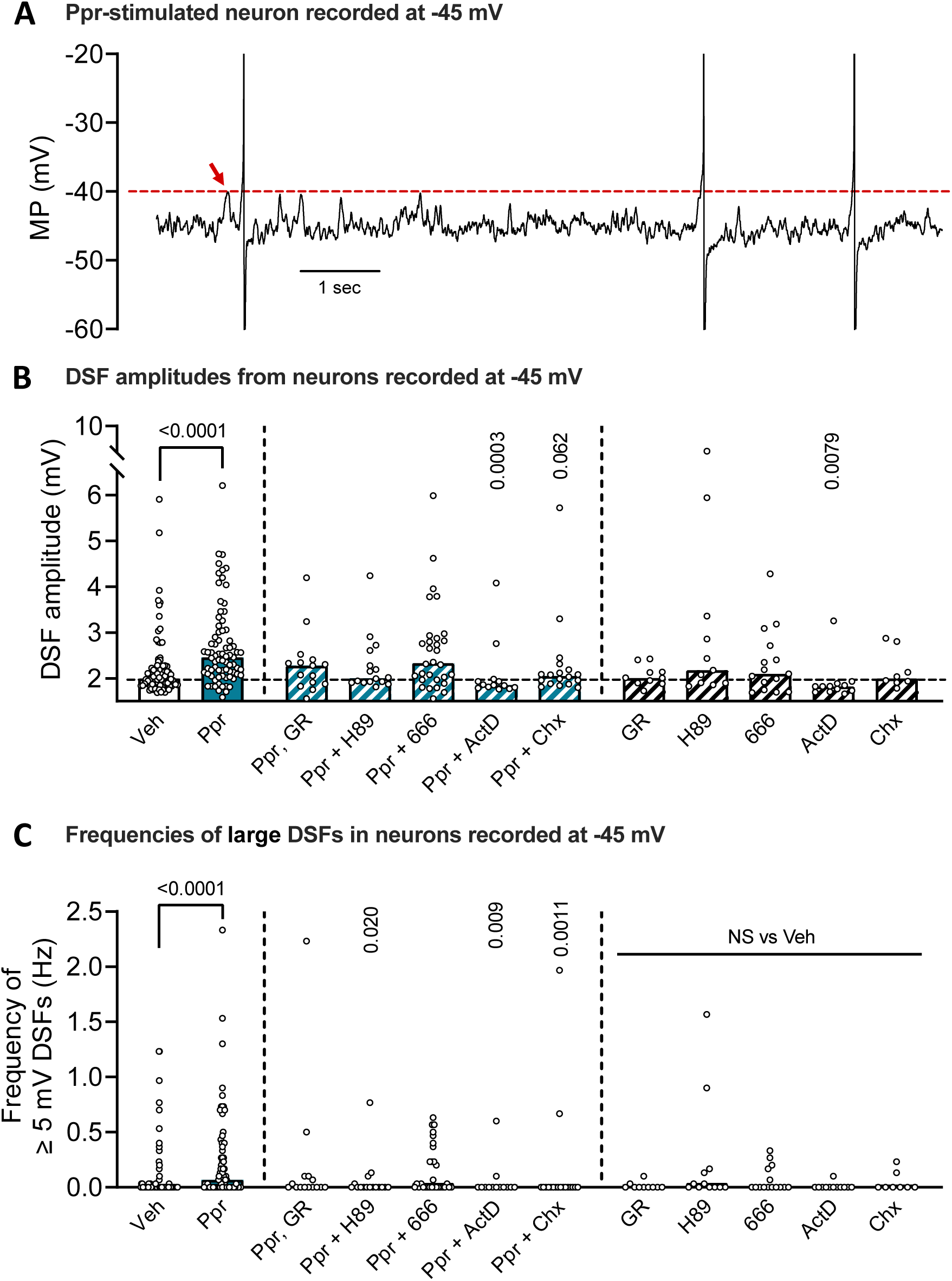
Ppr-induced LTH involves potentiation of DSFs. ***A***, magnification of the representative 10- second trace of a Ppr-treated sensory neuron recorded at -45 mV shown in Fig. 1D (right side). APs are clipped to show details of the fluctuations of membrane potential. Red arrow indicates the largest subthreshold DSF that was used as a conservative estimate of AP voltage threshold (represented by red dashed line) for subsequent analysis with FIBSI. ***B-C***, effect of Ppr and of each inhibitor tested on DSF amplitude (***B***) or the frequencies of large (≥ 5 mV) DSFs (***C***). Data are graphed as medians. Each open circle represents the mean DSF value from a single neuron. Comparisons of amplitudes or large DSF frequencies between Ppr and vehicle or each Ppr with inhibitor used Kruskal-Wallis followed by Dunn’s multiple comparisons tests, as were differences between vehicle and each inhibitor alone. *p* values are reported in each panel and *p*<0.05 was considered significant.

## 4. Discussion

This study has shown that a prototypical cellular signaling pathway previously shown to induce memory-related synaptic LTP can also induce long-lasting hyperexcitability in isolated nociceptor somata. Because nociceptor LTH is likely to promote hyperactivity that can drive central sensitization and ongoing pain (Walters et al., 2023; Walters, 2023), our findings provide insight into how persistent pain may be induced, and they raise the possibility that this cellular signaling pathway might also be involved in chronically maintaining LTH and hyperalgesic priming.

### 4.1. Prucalopride-induced LTH involves excitability alterations that both resemble and differ from those found in nociceptors in diverse pain models

Our investigation of dissociated nociceptors has demonstrated that temporary (6-hour) treatment with 5-HT4 receptor agonist prucalopride induces LTH lasting at least 1 day. This LTH exhibits similarities to and some differences from nociceptor hyperexcitability revealed by comparable current clamp test protocols in several pain models. A difference from the chronic hyperactivity induced by spinal cord injury (SCI) (Bedi et al., 2010; Odem et al., 2018; Berkey et al., 2020) and from the hyperactivity induced by *in vivo* cisplatin treatment (Laumet et al., 2020) is that the prucalopride- induced LTH did not manifest as overt hyperactivity (SA at RMP) – OA was only revealed during modest experimental depolarization to a normally subthreshold membrane potential (-45 mV). The combination of absent SA at RMP with enhanced OA evoked by modest depolarization is also observed acutely during 5-HT treatment *in vitro* (Odem et al., 2018; Lopez et al., 2021), and often for several weeks after plantar incision injury (Bavencoffe et al., 2024). Another difference from the SCI and cisplatin models is the lack of persistently depolarized RMP. On the other hand, distinctive similarities are found across these conditions, including an enhancement of DSFs and (in many cases) hyperpolarization of AP threshold. In general, SCI and cisplatin treatment induce stronger hyperexcitable alterations that produce continuing hyperactivity (SA at RMP), while the persistent effects of plantar incision, the acute effects of 5-HT treatment, and the long-lasting aftereffects of prucalopride treatment are expressed as weaker hyperexcitable alterations that remain latent until an event occurs to depolarize RMP and precipitate suprathreshold hyperactivity.

### 4.2. A fundamental, memory-related cell signaling pathway can induce nociceptor LTH

Our *in vitro* LTH induction protocol was modeled on pharmacological experiments using dissociated sensory and motor neurons from the marine snail, *Aplysia californica*, which first revealed a prototypical cell signaling pathway for inducing long-term enhancement of synaptic transmission related to memory formation. These classic studies used 5-HT treatment to show that stimulation of a cAMP-PKA-CREB-gene transcription-protein synthesis pathway for 1.5 hours was sufficient to enhance synaptic transmission 1 day later (Montarolo et al., 1986; Schacher et al., 1988; Dash et al., 1990; Casadio et al., 1999). Discovery of this long-term synaptic plasticity induction pathway associated with behaviorally expressed memory of noxious stimulation was recognized by a Nobel Prize in 2000 (Kandel, 2001). This general pathway was then implicated in memory-related late-phase LTP induction in the hippocampus and amygdala (Abraham et al., 1991; Frey et al., 1993; Impey et al., 1998; Huang et al., 2000; Barco et al., 2002) and spinal cord (Yang et al., 2004; Liu and Zhou, 2015). Induction signals for synaptic memory have since been found to be far more complex than the prototypical cAMP-PKA-CREB-transcription-translation pathway (Bito et al., 1996; Impey et al., 1998; Reul, 2014; Abbas et al., 2015). Nevertheless, our demonstration that nociceptor LTH can be induced by either a Gαs-coupled 5-HT receptor agonist (prucalopride) or forskolin and that prucalopride-induced LTH can be prevented by co-treatment with inhibitors of PKA, CREB, gene transcription, or protein synthesis is significant. It suggests that this pathway and related pathways having ancient transcription- and translation-dependent plasticity functions (Walters, 2023) are important both for inducing synaptic alterations subserving conventional memory formation and for inducing nociceptor memories of bodily injury expressed as continuing hyperexcitability . A role for this pathway in the induction of nociceptor memory is further supported by a report that *in vivo* injection of 8-bromo-cAMP into or close to a DRG induces hyperalgesic priming expressed as enhanced PGE2-elicited sensitization of paw withdrawal responses 1 week later (Ferrari et al., 2015). Especially noteworthy is that the priming induced by 8-bromo-cAMP was prevented by local application of the transcription inhibitor actinomycin D to the DRG or by intrathecal delivery of antisense oligodeoxynucleotides that knock down CREB (Ferrari et al., 2015). These observations encourage further *in vivo* investigation of the roles of the cAMP-PKA-CREB-gene transcription-protein synthesis pathway in the induction of nociceptor LTH and its contribution to persistent nociceptive sensitization. In addition, injury and inflammation *in vivo* should trigger the release of many sensitizing factors: not only molecules that stimulate Gαs-coupled receptors and cAMP signaling (e.g., 5-HT, prostaglandins), but also numerous cytokines and damage-associated signals that stimulate other pathways that potentially contribute to the induction of LTH and hyperalgesic priming.

### 4.3. How is induction of LTH by prucalopride in vitro related to nociceptor hyperexcitability induced in vivo by persistent pain conditions?

Hyperactivity in dissociated nociceptors (expressed as SA at RMP under whole-cell current clamp) has been observed from 3 days to at least 6-8 months after spinal cord injury (SCI) (Bedi et al., 2010; Odem et al., 2018), suggesting a very long-lasting memory of severe bodily injury.

Dissociated small DRG neurons (likely nociceptors) in rodents also exhibit hyperactivity expressed as SA and/or OA during moderate depolarization weeks after *in vivo* sciatic nerve injury (Study and Kral, 1996), weeks after plantar incision injury (Bavencoffe et al., 2024), and days after *in vivo* treatment with paclitaxel (Li et al., 2017) or cisplatin (Laumet et al., 2020). Moreover, SA in dissociated DRG neurons associated with previously reported pain was found in humans long after compression of spinal nerves or ganglia by tumors (North et al., 2019, 2022). In most of these studies, the hyperexcitability was present 1 day after dissociation, indicating that an intrinsic cellular memory of the painful condition lasts at least 1 day. Two general mechanisms -- not mutually exclusive -- could explain nociceptor LTH persisting for weeks, months, or longer *in vivo*. One is analogous to synaptic memory models that posit a transient induction event, often assumed to require activation of immediate early genes and an ensuing cascade of transcriptional and translational effects, that trigger enduring neuronal alterations (analogous to transcriptional and translational cascades important for development) (Kandel, 2001). The other possibility is that there is a continuing reinduction of shorter- lasting neuronal alterations by recurring extrinsic stimulation, e.g., recurrent or continuing exposure of a nociceptor to inflammatory mediators such as 5-HT long after an initiating injury. Both types of persistence mechanisms are proposed to operate in conventional memory, where a memory that was 1) initially consolidated (formed) by transcription- and translation-dependent mechanisms can be 2) updated and extended by repeated reconsolidation via transcriptional and translational mechanisms each time the memory is reactivated (Dudai, 2012; Alberini and Kandel, 2014; Bonin and De Koninck, 2015). The present results provide proof-of-principle for the first possibility: that early, temporary cAMP-induced transcriptional and translational mechanisms can induce nociceptor LTH that might last for long periods.

At the same time, our results are consistent with the second, complementary possibility: that nociceptor LTH is “reconsolidated” by continuing re-exposure to extrinsic sensitizing signals. For example, plasma levels of PGE2, which induces acute nociceptor hyperexcitability via cAMP signaling (Taiwo et al., 1989; England et al., 1996; Wang et al., 2007), have been found to increase in a painful chronic arthritis model (Feuerherm et al., 2019). In some conditions, circulating sensitizing signals that might ebb and flow over time may repeatedly re-induce LTH in peripheral terminals and/or the cell body (which lacks an effective vascular permeability barrier (Abram et al., 2006; Jimenez-Andrade et al., 2008)) so that the LTH persists much longer than it would after only a single, temporary exposure. Injured tissue is at risk for reinjury and is usually sensitized. Sensitization increases the chance that subsequent noxious stimulation, or stimulation that would ordinarily be innocuous but poses a threat to weakened tissue, causes the further release of sensitizing signals (e.g., 5-HT, PGE2) that can reactivate transcription- and translation-dependent pathways to prolong nociceptor LTH and other protective sensitizing effects while healing progresses. This adaptive perspective suggests that inflammatory challenges, such as reinjury or experimental PGE2 injection, coming after events that induce hyperalgesic priming may re-activate the cAMP-PKA-transcription-translation pathway and other cellular memory induction pathways to extend protective sensitization during periods of increased tissue vulnerability. A third (also complementary) possibility is that positive feedback among cell signaling loops independently of transcriptional and translational mechanisms helps to maintain persistent hyperexcitability, as suggested by our findings of positive feedback among cAMP signaling, ERK signaling, Ca^2+^ signaling, and depolarization long after SCI (Garza Carbajal et al., 2020; 2024). It is plausible that all three general mechanisms contribute to persistent nociceptor hyperexcitability and that their individual contributions depend upon the nature and severity of the initiating biological insult and the degree of subsequent exposure to stressors of weakened tissue.

## Disclosures

This work was supported by grants from the National Institutes of Health: NS091759 (Carmen W. Dessauer and Edgar T. Walters), NS111521 (Edgar T. Walters and Michael X. Zhu), as well as the Fondren Chair in Cellular Signaling (Edgar T. Walters) The authors declare no conflicts of interest.

## Notes

### Competing Interest Statement

The authors have declared no competing interest.

## References

Abbas AK, Villers A, Ris L (2015) Temporal phases of long-term potentiation (LTP): myth or fact. Rev Neurosci, 26:507–546.

Abraham WC, Dragunow M, Tate WP (1991) The role of immediate early genes in the stabilization of long-term potentiation. Mol Neurobiol, 5:297–314.

Abram SE, Yi J, Fuchs A, Hogan QH (2006) Permeability of injured and intact peripheral nerves and dorsal root ganglia. Anesthesiology, 105:146–153.

Alberini CM, Kandel ER (2014) The regulation of transcription in memory consolidation. Cold Spring Harb Perspect Biol, 7:a021741.

Barco A, Alarcon JM, Kandel ER (2002) Expression of constitutively active CREB protein facilitates the late phase of long-term potentiation by enhancing synaptic capture. Cell, 108:689–703.

Basbaum AI, Bautista DM, Scherrer G, Julius D (2009) Cellular and molecular mechanisms of pain. Cell, 139:267–284.

Bavencoffe A, Spence EA, Zhu MY, Garza-Carbajal A, Chu KE, Bloom OE, Dessauer CW, Walters ET (2022) Macrophage Migration Inhibitory Factor (MIF) Makes Complex Contributions to Pain- Related Hyperactivity of Nociceptors after Spinal Cord Injury. J Neurosci, 42:5463–5480.

Bavencoffe AG, Lopez ER, Johnson KN, Tian J, Gorgun FM, Shen BQ, Zhu MX, Dessauer CW, Walters ET (2024) Widespread latent hyperactivity of nociceptors outlasts enhanced avoidance behavior following incision injury. bioRxiv, 2024.01.30.578108.

Bedi SS, Yang Q, Crook RJ, Du J, Wu Z, Fishman HM, Grill RJ, Carlton SM, Walters ET (2010) Chronic spontaneous activity generated in the somata of primary nociceptors is associated with pain-related behavior after spinal cord injury. J Neurosci, 30:14870–14882.

Berkey SC, Herrera JJ, Odem MA, Rahman S, Cheruvu SS, Cheng X, Walters ET, Dessauer CW, Bavencoffe AG (2020) EPAC1 and EPAC2 promote nociceptor hyperactivity associated with chronic pain after spinal cord injury. Neurobiol Pain, 7:100040.

Bito H, Deisseroth K, Tsien RW (1996) CREB phosphorylation and dephosphorylation: a Ca(2+)- and stimulus duration-dependent switch for hippocampal gene expression. Cell, 87:1203–1214.

Bonin RP, De Koninck Y (2015) Reconsolidation and the regulation of plasticity: moving beyond memory. Trends Neurosci, 38:336–344.

Casadio A, Martin KC, Giustetto M, Zhu H, Chen M, Bartsch D, Bailey CH, Kandel ER (1999) A transient, neuron-wide form of CREB-mediated long-term facilitation can be stabilized at specific synapses by local protein synthesis. Cell, 99:221–237.

Cudmore RH, Turrigiano GG (2004) Long-term potentiation of intrinsic excitability in LV visual cortical neurons. J Neurophysiol, 92:341–348.

Dash PK, Hochner B, Kandel ER (1990) Injection of the cAMP-responsive element into the nucleus of Aplysia sensory neurons blocks long-term facilitation. Nature, 345:718–721.

Davis RL (2023) Learning and memory using Drosophila melanogaster: a focus on advances made in the fifth decade of research. Genetics, 224:iyad085.

Dudai Y (2012) The restless engram: consolidations never end. Annu Rev Neurosci, 35:227–247.

England S, Bevan S, Docherty RJ (1996) PGE2 modulates the tetrodotoxin-resistant sodium current in neonatal rat dorsal root ganglion neurones via the cyclic AMP-protein kinase A cascade. J Physiol, 495:429–440.

Ferrari LF, Araldi D, Levine JD (2015) Distinct terminal and cell body mechanisms in the nociceptor mediate hyperalgesic priming. J Neurosci, 35:6107–6116.

Feuerherm AJ, Dennis EA, Johansen B (2019) Cytosolic group IVA phospholipase A2 inhibitors, AVX001 and AVX002, ameliorate collagen-induced arthritis. Arthritis Res Ther, 21:29.

Finnerup NB, Kuner R, Jensen TS (2021) Neuropathic Pain: From Mechanisms to Treatment. Physiol Rev, 101:259–301.

Frey U, Huang YY, Kandel ER (1993) Effects of cAMP simulate a late stage of LTP in hippocampal CA1 neurons. Science, 260:1661–1664.

Gale JR, Gedeon JY, Donnelly CJ, Gold MS (2022) Local translation in primary afferents and its contribution to pain. Pain, 163:2302–2314.

Garza-Carbajal A, Bavencoffe A, Walters ET, Dessauer CW (2020) Depolarization-Dependent C-Raf Signaling Promotes Hyperexcitability and Reduces Opioid Sensitivity of Isolated Nociceptors after Spinal Cord Injury. J Neurosci, 40:6522–6535.

Garza-Carbajal A, Bavencoffe A, Herrera JJ, Johnson, KN, Walters ET, Dessauer CW (2024) Mechanism of Gabapentinoid Potentiation of Opioid Effects on cAMP Signaling in Neuropathic Pain. Proc Natl Acad Sci USA, in press.

Ghosh K, Pan HL (2022) Epigenetic Mechanisms of Neural Plasticity in Chronic Neuropathic Pain. ACS Chem Neurosci, 13:432–441.

Huang EP, Stevens CF (1998) The matter of mind: molecular control of memory. Essays Biochem, 33:165–178.

Huang YY, Martin KC, Kandel ER (2000) Both protein kinase A and mitogen-activated protein kinase are required in the amygdala for the macromolecular synthesis-dependent late phase of long-term potentiation. J Neurosci, 20:6317–6325.

Impey S, Obrietan K, Wong ST, Poser S, Yano S, Wayman G, Deloulme JC, Chan G, Storm DR (1998) Cross talk between ERK and PKA is required for Ca2+ stimulation of CREB-dependent transcription and ERK nuclear translocation. Neuron, 21:869–883.

Ji RR, Kohno T, Moore KA, Woolf CJ (2003) Central sensitization and LTP: do pain and memory share similar mechanisms. Trends Neurosci, 26:696–705.

Jimenez-Andrade JM, Herrera MB, Ghilardi JR, Vardanyan M, Melemedjian OK, Mantyh PW (2008) Vascularization of the dorsal root ganglia and peripheral nerve of the mouse: implications for chemical-induced peripheral sensory neuropathies. Mol Pain, 4:10.

Kandel ER (2001) The molecular biology of memory storage: a dialogue between genes and synapses. Science, 294:1030–1038.

Kandel ER (2012) The molecular biology of memory: cAMP, PKA, CRE, CREB-1, CREB-2, and CPEB. Mol Brain, 5:14.

Laumet G, Bavencoffe A, Edralin JD, Huo XJ, Walters ET, Dantzer R, Heijnen CJ, Kavelaars A (2020) Interleukin-10 resolves pain hypersensitivity induced by cisplatin by reversing sensory neuron hyperexcitability. Pain, 161:2344–2352.

Li XH, Miao HH, Zhuo M (2019) NMDA Receptor Dependent Long-term Potentiation in Chronic Pain. Neurochem Res, 44:531–538.

Li Y, Tatsui CE, Rhines LD, North RY, Harrison DS, Cassidy RM, Johansson CA, Kosturakis AK, Edwards DD, Zhang H, Dougherty PM (2017) Dorsal root ganglion neurons become hyperexcitable and increase expression of voltage-gated T-type calcium channels (Cav3.2) in paclitaxel-induced peripheral neuropathy. Pain, 158:417–429.

Liao X, Gunstream JD, Lewin MR, Ambron RT, Walters ET (1999) Activation of protein kinase A contributes to the expression but not the induction of long-term hyperexcitability caused by axotomy of Aplysia sensory neurons. J Neurosci, 19:1247–1256.

Liu XG, Zhou LJ (2015) Long-term potentiation at spinal C-fiber synapses: a target for pathological pain. Curr Pharm Des, 21:895–905.

Lopez ER, Carbajal AG, Tian JB, Bavencoffe A, Zhu MX, Dessauer CW, Walters ET (2021) Serotonin enhances depolarizing spontaneous fluctuations, excitability, and ongoing activity in isolated rat DRG neurons via 5-HT_4_ receptors and cAMP-dependent mechanisms. Neuropharmacology, 184:108408.

Mihail SM, Wangzhou A, Kunjilwar KK, Moy JK, Dussor G, Walters ET, Price TJ (2019) MNK-eIF4E signalling is a highly conserved mechanism for sensory neuron axonal plasticity: evidence from Aplysia californica. Philos Trans R Soc Lond B Biol Sci, 374:20190289.

Montarolo PG, Goelet P, Castellucci VF, Morgan J, Kandel ER, Schacher S (1986) A critical period for macromolecular synthesis in long-term heterosynaptic facilitation in Aplysia. Science, 234:1249– 1254.

Nguyen PV, Woo NH (2003) Regulation of hippocampal synaptic plasticity by cyclic AMP-dependent protein kinases. Prog Neurobiol, 71:401–437.

North RY, Li Y, Ray P, Rhines LD, Tatsui CE, Rao G, Johansson CA, Zhang H, Kim YH, Zhang B, Dussor G, Kim TH, Price TJ, Dougherty PM (2019) Electrophysiological and transcriptomic correlates of neuropathic pain in human dorsal root ganglion neurons. Brain, 142:1215–1226.

North RY, Odem MA, Li Y, Tatsui CE, Cassidy RM, Dougherty PM, Walters ET (2022) Electrophysiological alterations driving pain-associated spontaneous activity in human sensory neuron somata parallel alterations described in spontaneously active rodent nociceptors. J Pain, S1526–5900(22)00047.

Odem MA, Bavencoffe AG, Cassidy RM, Lopez ER, Tian J, Dessauer CW, Walters ET (2018) Isolated nociceptors reveal multiple specializations for generating irregular ongoing activity associated with ongoing pain. Pain, 159:2347–2362.

Price TJ, Ray PR (2019) Recent advances toward understanding the mysteries of the acute to chronic pain transition. Curr Opin Physiol, 11:42–50.

Reichling DB, Levine JD (2009) Critical role of nociceptor plasticity in chronic pain. Trends Neurosci, 32:611–618.

Reul JM (2014) Making memories of stressful events: a journey along epigenetic, gene transcription, and signaling pathways. Front Psychiatry, 5:5.

Sandkühler J (2007) Understanding LTP in pain pathways. Mol Pain, 3:9.

Schacher S, Castellucci VF, Kandel ER (1988) cAMP evokes long-term facilitation in Aplysia sensory neurons that requires new protein synthesis. Science, 240:1667–1669.

Scholz KP, Byrne JH (1988) Intracellular injection of cAMP induces a long-term reduction of neuronal K+ currents. Science, 240:1664–1666.

Silva AJ, Kogan JH, Frankland PW, Kida S (1998) CREB and memory. Annu Rev Neurosci, 21:127– 148.

Study RE, Kral MG (1996) Spontaneous action potential activity in isolated dorsal root ganglion neurons from rats with a painful neuropathy. Pain, 65:235–242.

Taiwo YO, Bjerknes LK, Goetzl EJ, Levine JD (1989) Mediation of primary afferent peripheral hyperalgesia by the cAMP second messenger system. Neuroscience, 32:577–580.

Tian J, Bavencoffe AG, Zhu MX, Walters ET (2024) Readiness of nociceptor cell bodies to generate spontaneous activity results from background activity of diverse ion channels and high input resistance. Pain, 165:893–907.

Walters ET (1991) A Functional, Cellular, and Evolutionary Model of Nociceptive Plasticity in Aplysia. Biol Bull, 180:241–251.

Walters ET (2023) Exaptation and Evolutionary Adaptation in Nociceptor Mechanisms Driving Persistent Pain. Brain Behav Evol, 98:314–330.

Walters ET, Ambron RT (1995) Long-term alterations induced by injury and by 5-HT in Aplysia sensory neurons: convergent pathways and common signals. Trends Neurosci, 18:137–142.

Walters ET, Crook RJ, Neely GG, Price TJ, Smith ESJ (2023) Persistent nociceptor hyperactivity as a painful evolutionary adaptation. Trends Neurosci, 46:211–227.

Wang C, Li GW, Huang LY (2007) Prostaglandin E2 potentiation of P2X3 receptor mediated currents in dorsal root ganglion neurons. Mol Pain, 3:22.

Weragoda RM, Ferrer E, Walters ET (2004) Memory-like alterations in Aplysia axons after nerve injury or localized depolarization. J Neurosci, 24:10393–10401.

Weragoda RM, Walters ET (2007) Serotonin induces memory-like, rapamycin-sensitive hyperexcitability in sensory axons of aplysia that contributes to injury responses. J Neurophysiol, 98:1231–1239.

Woolf CJ, Walters ET (1991) Common patterns of plasticity contributing to nociceptive sensitization in mammals and Aplysia. Trends Neurosci, 14:74–78.

Wu Z, Yang Q, Crook RJ, O’Neil RG, Walters ET (2013) TRPV1 channels make major contributions to behavioral hypersensitivity and spontaneous activity in nociceptors after spinal cord injury. Pain, 154:2130–2141.

Yang HW, Hu XD, Zhang HM, Xin WJ, Li MT, Zhang T, Zhou LJ, Liu XG (2004) Roles of CaMKII, PKA, and PKC in the induction and maintenance of LTP of C-fiber-evoked field potentials in rat spinal dorsal horn. J Neurophysiol, 91:1122–1133.

